# A rhomboid protease, EhROM1, regulates the submembrane distribution and size of the Gal/GalNAc lectin subunits in the human protozoan parasite, *Entamoeba histolytica*

**DOI:** 10.1101/693044

**Authors:** Brenda H. Welter, Lesly A. Temesvari

**Author notes:** Corresponding author, Lesly A. Temesvari (LAT).

## Abstract

*Entamoeba histolytica* is a food- and waterborne parasite that is the causative agent of amebic dysentery and amoebic liver abscesses. Adhesion is one of the most important virulence functions as it facilitates motility, colonization of host, destruction of host tissue, and uptake of nutrients by the parasite. One well-characterized parasite cell surface adhesin is the Gal/GalNAc lectin, which binds to galactose or *N*-acetylgalactosamine residues on host components and is composed of heavy (Hgl), intermediate (Igl), and light (Lgl) subunits. Igl has been shown to be constitutively localized to lipid rafts (cholesterol-rich membrane domains), whereas Hgl and Lgl transiently associate with rafts. When all three subunits are localized to rafts there is an increase in galactose-sensitive adhesion. Thus, submembrane location may regulate the function of this adhesion. Rhomboid proteases are a conserved family of intramembrane proteases that also participate in the regulation of parasite-host interactions. In *E. histolytica*, one rhomboid protease, EhROM1, cleaves Hgl as a substrate, and knockdown of its expression inhibits parasite-host interactions. Since rhomboid proteases are found within membranes, it is not surprising that lipid composition regulates their activity and enzyme-substrate binding. Given the importance of the lipid environment for both rhomboid proteases and the Gal/GalNAc lectin, we sought to gain insight into the relationship between rhomboid proteases and submembrane location of the lectin in *E. histolytica*. We demonstrated that EhROM1, itself, is enriched in rafts. Reducing rhomboid protease activity, either pharmacologically or genetically, correlated with an enrichment of Hgl and Lgl in rafts. In a mutant cell line with reduced EhROM1 expression, there was also a significant augmentation of the level of all three Gal/GalNAc subunits on the cell surface and an increase in the molecular weight of Hgl and Lgl. Overall, the study provides insight into the molecular mechanisms governing parasite-host adhesion for this pathogen.

## Introduction

*Entamoeba histolytica* is an intestinal parasite that is the causative agent of amebic dysentery and amoebic liver abscesses (reviewed in [1]). It is transmitted via the cyst form of the pathogen in fecally-contaminated food and water, making it prevalent in the developing world where sanitation practices are substandard. In 2015, it was estimated that 2.4 billion people still lack access to improved sanitation facilities and nearly 1 billion people carry out open defecation practices [2]. Thus, there is considerable risk for the transmission of *E. histolytica*.

After ingestion, the cyst exits the stomach and enters the small intestine, where unknown triggers cause excystation. The emerging amoeboid trophozoites travel down the digestive system until they reach the large intestine, where infection is established. In the large intestine the trophozoites feed on bacteria and host cell material by endocytosis, and divide by binary fission. Trophozoites can also invade the colonic epithelial lining and cause extra-intestinal complications of infection, including liver abscess. During colonization and invasion of the host, trophozoites adhere to numerous host-derived and host flora-derived ligands, including epithelial cells [3], red blood cells (RBCs) [4,5], extracellular matrix components (e.g., collagen [4,6] and fibronectin [7]), intestinal flora [8,9], colonic mucins [8–10], and leukocytes [11,12]. Since adhesion is one of the first steps in host colonization, and facilitates uptake of nutrients by endocytosis, it may be one of the most important of virulence functions for this parasite.

Several parasite cell surface adhesins have been described (reviewed in [13]); however, the best characterized is the heterotrimeric protein complex known as the galactose/N-acetylgalactosamine lectin (Gal/GalNAc lectin). This complex binds to galactose and N-acetylgalactosamine residues on host ligands and is composed of heavy (Hgl), light (Lgl), and intermediate (Igl) subunits. Hgl is a transmembrane protein that possesses an exoplasmic carbohydrate binding domain and is disulfide-linked to glycophosphatidylinositol (GPI)-anchored Lgl. The Hgl-Lgl heterodimer non-covalently associates with GPI-anchored Igl. Both Hgl and Igl share sequence homology with β integrins [14–17], suggesting that they may also play a role in signaling. In particular, Hgl is thought to facilitate signaling and adhesion not only by binding extracellular ligands, but also by interacting with proteins in the intracellular space of the parasite, via its cytoplasmic tail [15].

Functional regulation of the Gal/GalNAc lectin is not well understood; however, it has been shown that lipid rafts may play a role [4,6,18,19]. Lipid rafts are tightly packed cholesterol- and sphingolipid-rich membrane microdomains that serve as compartments within which signaling proteins interact. It was previously shown that the majority of Igl is found in raft-like membranes, whereas the majority of Hgl and Lgl is found in non-raft membrane [18]. However, exposing *E. histolytica* cells to a source of cholesterol [19] or several host ligands, including RBCs [4] or collagen [6], results in enrichment of Hgl and Lgl in rafts and thus, co-compartmentalisation of all three subunits. Colocalization of the subunits in rafts is accompanied by an increase in the ability of the amoebae to adhere to host components in a galactose-specific manner [19]. Removal of cholesterol disrupts lipid rafts and inhibits the adhesion of *E. histolytica* trophozoites to host cells [18] and collagen [6]. Together, these data suggest that there is a correlation between sub-membrane location and function of the Gal/GalNAc lectin, and that lipid rafts may serve as a platform for the assembly, modification, and/or functional regulation of proteins involved in parasite-host interaction.

Cells must also possess mechanisms to modulate or dismantle adhesion junctions. Rhomboid proteases are a family of intramembrane proteases that participate in a wide variety of cellular functions including cell signaling, mitochondrial homeostasis, quorum sensing, protein translocation across membranes and the regulation of adhesion junctions (reviewed in [20]). They are conserved from bacteria to mammals and their role in regulating parasite-host interactions (reviewed in [21]) is established in *Toxoplasma* [22–26], *Plasmodium* [27–29], *Trichomonas* [30], and *Entamoeba* [31–33].

In particular, for *E. histolytica*, Hgl is thought to be a substrate of the *E. histolytica* rhomboid protease, EhROM1, because it possesses a canonical rhomboid protease-binding sequence and it can be cleaved by EhROM1 when they are co-expressed in a mammalian cell system [31]. Knocking down expression of EhROM1 using an epigenetic silencing approach results in reduced adhesion to host cells and reduced erythrophagocytosis [32]. Overexpression of a dominant negative catalytic mutant of EhROM1 also causes defects in host cell binding [33]. Finally, overexpression of the catalytic mutant or knocking down expression, using an RNAi-based method, gives rise to mutant cells that are less cytotoxic, hemolytic, and motile than control cells [33]. Together, these observations support the role of EhROM1 in parasite-host interactions.

Since rhomboid proteases have an intramembrane position, a logical conjecture is that lipid composition regulates activity and compartmentalization regulates enzyme-substrate contact. In support of this, the activity of both prokaryotic and eukaryotic rhomboid proteases can be influenced by membrane composition *in vitro* [34] and pharmacological perturbation of cellular membranes *in vivo* can alter the activity of at least one rhomboid protease, human RHBDL4 [35].

Given the importance of compartmentalization for both rhomboid proteases and the Gal/GalNAc lectin, we sought to gain insight into the relationship between rhomboid protease activity and submembrane location of the lectin in *E. histolytica*. We demonstrate that EhROM1 is localized to lipid rafts and loss of rhomboid protease expression correlates with an enrichment of the Gal/GalNAc lectin at the cell surface and in lipid rafts. We also show that the molecular weights of Hgl and Igl are increased in the absence of EhROM1 activity, supporting the notion that they are substrates of this protease. Overall, the study provides insight into the molecular mechanisms governing parasite-host adhesion for this pathogen.

## Methods

### Strains and culture conditions

The generation of an *Entamoeba histolytica* cell line with RNAi-mediated reduced expression of EhROM1 is described elsewhere [33], and was generously provided by Dr. Upinder Singh (Division of Infectious Diseases, Dept. of Internal Medicine, Dept. of Microbiology and Immunology, Stanford University School of Medicine, Stanford, CA, USA). Both mutant and wildtype *E. histolytica* trophozoites (strain HM-1:IMSS) were cultured axenically in TYI-S-33 media [36] in 15 ml glass screw cap tubes.

### Pharmacological inhibition of rhomboid protease activity

To inhibit rhomboid protease activity, parasites (3.5 x 10^6^ cells/ml) were treated with 100 μM 3,4-dichloroisocoumarin (DCI) (Sigma-Aldrich, St. Louis, MO). DCI was dissolved in dimethyl sulfoxide (DMSO) and applied to the parasites for 2 h at 37°C. Control parasites were treated with DMSO alone.

### Lipid raft isolation and characterization by western blot analysis

Isolation and characterization of lipid rafts were carried out as previously described [18]. Briefly, 3 × 10^6^ *Entamoeba* cells were incubated at 4°C for 30 min in ice cold Buffer 1 containing protease inhibitors (40 mM sodium pyrophosphate, 0.4 mM dithiothreitol, 0.1 mg of phenylmethylsulfonyl fluoride/ml, 2 mM EDTA, 1 mM EGTA, 3 mM sodium azide, 10 mM Tris-HCl [pH 7.6]) and 0.5% (v/v) Triton X-100. The cells were then collected by centrifugation (14,500 × g, 5 min) at 4°C. The Triton-soluble supernatant (TSS) was removed, and the Triton-insoluble pellet (TIP) was resuspended in 80% (wt/vol) sucrose in Buffer 1. A noncontinuous sucrose gradient was generated by using equal volumes of 80 (containing the TIP), 50, 30, and 10% (wt/vol) sucrose solutions in Buffer 1. Samples were then centrifuged in a Beckman Coulter (Indianapolis, IN) Optima™ MAX-XP ultracentrifuge (125,000 × g for 16 h at 4°C). After centrifugation, the gradient was fractionated into 20 equal volumes (140 μl/fraction). The proteins in each sample were precipitated by the addition of trichloroacetic acid (TCA) as described elsewhere [37]. The precipitated proteins were resuspended in double-distilled H_2_O and mixed with 4× LDS buffer (Invitrogen, Carlsbad, CA) and 2-mercaptoethanol (10% [vol/vol] final concentration).

TCA-precipitated proteins were analyzed by SDS-PAGE and western blot analysis as previously described [4,18,19]. Primary antibodies for western blot characterization of the sucrose gradient fractions included a mixture of monoclonal α-Lgl antibodies (3C2, IC8, IA9, and ID4) (1:4,000 dilution), polyclonal α-Hgl antibodies (1:5,000 dilution) [38], a mixture of monoclonal α-Igl antibodies (3G5-A3-G3, 5H1-F11-D11, and 4G2-D8-H1) (1:4,000 dilution), or polyclonal α-EhROM1 antibodies (1:50 dilution). Monoclonal antibodies recognizing Igl and Lgl were generously provided by Dr. William A. Petri, Jr. (Division of Infectious Diseases and International Health, University of Virginia, Charlottesville, VA, USA). Polyclonal antibodies recognizing EhROM1 were generously provided by Dr. Upinder Singh (Division of Infectious Diseases, Dept. of Internal Medicine, Dept. of Microbiology and Immunology, Stanford University School of Medicine, Stanford, CA, USA). Bethesda, MD).

Commercial HRP-conjugated polyclonal goat anti-rabbit or goat anti-mouse secondary antibodies (MP Biomedicals, Solon, OH) were utilized at 2 μg/ml. Western blots were analyzed by densitometry using ImageJ software (version 1.51; U.S. National Institutes of Health).

### Cell surface biotinylation

Control and T-EhROM1-s cells (6.0 × 10^5^) were surface biotinylated and purified by avidin affinity chromatography using the Pierce Cell Surface Protein Isolation Kit (Pierce Biotechnology, Rockford, IL, USA) according to manufacturer’s specifications. Flow-through fractions and biotinylated surface proteins that were captured by avidin affinity chromatography were resolved by SDS-PAGE and analyzed for Hgl, Lgl, Igl and actin by western blotting and densitometry as described above.

### Immunoprecipitation

Triton-insoluble membrane was isolated from wildtype and T-EhROM1-s cells as described above and fractionated on sucrose gradients. Fractions 11-12 (lipid rafts), 18-19 (actin-rich membrane) and TSS were used for immunoprecipitation assays. Combined fractions 11 and 12, 18 and 19 or TSS were pre-cleared by incubation with 1 × 10^7^ Dynabeads magnetic beads conjugated to sheep α-mouse IgG (Invitrogen Dynal AS, Oslo Norway) at 4°C for 2 h. The beads were collected in a microfuge tube magnetic separation stand (Promega, Madison, WI) and discarded. Protease inhibitors (as described above) and a mixture of monoclonal α-Hgl antibodies (1G7:3F4; ratio 3:1 to give a final antibody concentration of 0.01 μg/μl) were added to the pre-cleared fractions. The fractions were incubated for 1 h at room temperature with constant rotation. Monoclonal antibodies recognizing Hgl were generously provided by Dr. William A. Petri, Jr. (Division of Infectious Diseases and International Health, University of Virginia, Charlottesville, VA, USA). 1.5 × 10^7^ magnetic beads were then added to the samples, which were incubated overnight at 4 °C with constant rotation. Beads were collected in the magnetic separation stand, washed 6X with PBS, and boiled for 4 min in 4× LDS buffer and 2-mercaptoethanol (10% [vol/vol] final concentration). Samples were stored at −80 °C until analyzed by western blotting as described above.

### Statistical analyses

All values are given as means ± standard deviations (SD). Statistical analyses were performed using GraphPad Prism V.6.05 with a one-way analysis of variance (ANOVA) and a Tukey-Kramer multiple comparison test. *P* values of less than 0.05 were considered statistically significant. *P* values less than 0.01 were considered highly statistically significant.

## Results

### Inhibition of rhomboid proteases alters the sub-membrane location of the Gal/GalNAc subunits

Intramembrane protease-substrate interactions may be regulated by spatial segregation of either the enzyme or target protein in sub-membrane domains, such as lipid rafts [39]. Furthermore, lipid composition may influence intramembrane protease activity [34,35]. Thus, a logical prediction is that intramembrane proteolysis by rhomboid proteases may regulate the sub-membrane distribution of their substrates.

To determine if rhomboid protease activity influences the submembrane distribution of the subunits of the Gal/GalNAc lectin, we characterized the lipid rafts, as previously described [18], in trophozoites exposed to DCI, a compound which inhibits rhomboid proteases by alkylating an active-site histidine [34]. The lipid composition of rafts confers detergent resistance. Therefore, purification of lipid rafts was initiated by extraction with cold Triton X-100. This resulted in the isolation of detergent-resistant membrane (DRM), which consists of both lipid raft and actin-rich membrane. Since the buoyant density of lipid rafts is less than that of actin-rich membrane, these two membrane domains were further separated by sucrose density gradient centrifugation.

Western blot analysis of sucrose gradient fractions revealed that the majority of Igl was found in a low-density region (fractions 9 to 14) (Fig. 1). Previously, these fractions were shown to possess the highest levels of cholesterol, and thus are considered lipid rafts [18]. The localization of Igl to these low-density rafts is consistent with previous reports [4,18,19]. In control cells, the majority of Hgl and Lgl was associated with less buoyant, actin-rich fractions (fractions 17 to 20) (Fig. 1). However, after exposure to DCI, there was an increase in the proportion of Hgl and Lgl that was localized to lipid raft fractions (fractions 9 to 14), whereas the submembrane distribution of Igl remained unchanged (Fig. 1). Thus, inhibition of rhomboid protease activity correlates with an enrichment of Hgl and Lgl in lipid rafts in *E. histolytica*. Although, Lgl has not been identified as a EhROM1 substrate, the DCI-induced altered distribution of this subunit may be the result of its covalent connection to Hgl.

**Fig. 1.**
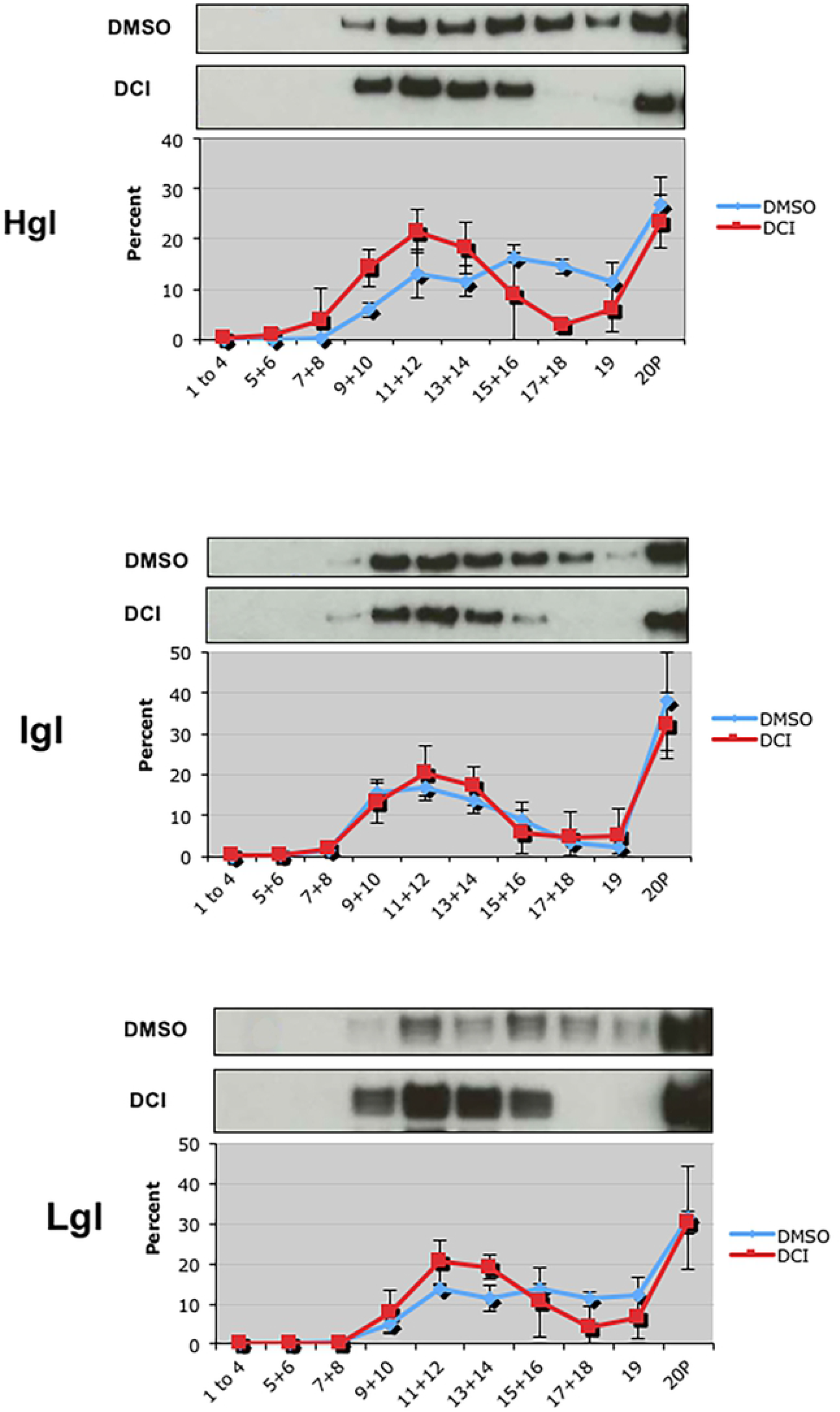
Inhibition of rhomboid protease activity with 3,4-dichloroisocoumarin (DCI) induces enrichment of the heavy (Hgl) and light (Lgl) subunits of the Gal/GalNAc lectins in lipid rafts. *E. histolytica* trophozoites were treated with a 100 μM DCI or diluent (DMSO) for 2 h. Triton-insoluble membranes were isolated and resolved by sucrose gradient density centrifugation. Nineteen fractions and the non-buoyant pellet (20P) were collected and subjected to western blot analyses using antibodies specific for the heavy subunit (Hgl), the intermediate subunit (Igl) or the light subunit (Lgl). Fractions 1 through 4, and pairs of fractions thereafter, were combined prior to analysis. Mean values of densitometric scans (n ≥ 2), reported as a percentage of total detergent-resistant membrane (DRM)-associated protein, are shown for each subunit. Representative western blots are shown above each panel. In both control (DMSO) and treated cells (DCI) Igl remained localized to central fractions (9-14) previously identified as lipid rafts [18]. In untreated cells (blue line) Hgl and Lgl predominantly localized to high density actin-rich fractions 15-19). After DCI-treatment (red line), Hgl and Lgl were enriched in low density lipid rafts.

Since chemical inhibitors may have off-target effects, we also used a genetic approach to corroborate the findings made after DCI treatment. Specifically, we isolated lipid rafts and characterized the sub-membrane distribution of the lectin subunits in a cell line (T-EhROM1-s) with reduced expression of EhROM1 [33]. Like DCI-treated cells, T-EhROM1-s cells exhibited augmented levels of Hgl and Lgl in lipid rafts (Fig. 2). These data support the notion that there is an inverse relationship between the level of rhomboid protease activity and the level of Hgl and Lgl in lipid rafts.

**Fig. 2.**
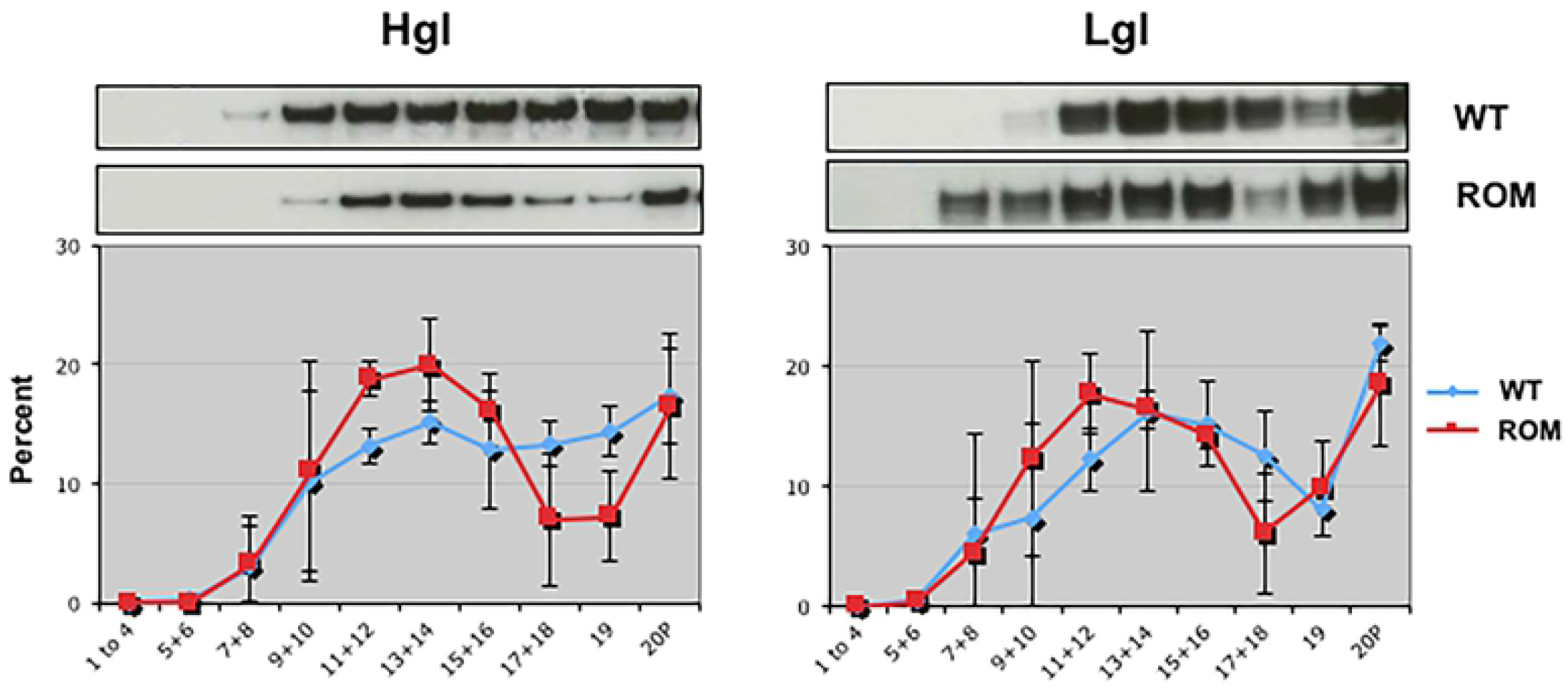
Reduced expression of EhROM1 correlates with enrichment of the heavy (Hgl) and light (Lgl) subunits of the Gal/GalNAc lectins in lipid rafts. Triton-insoluble membranes were isolated from wildtype cells (WT) or T-EhROM1-s (ROM) cells and resolved by sucrose gradient density centrifugation. Nineteen fractions and the non-buoyant pellet (20P) were collected and subjected to western blot analyses using antibodies specific for the heavy subunit (Hgl) or the light subunit (Lgl). Fractions 1 through 4, and pairs of fractions thereafter, were combined prior to analysis. Mean values of densitometric scans (n ≥ 3), reported as a percentage of total detergent-resistant membrane (DRM)-associated protein, are shown for each subunit. Representative western blots are shown above each panel. In T-EhROM1-s cells (red line), both Hgl and Lgl are enriched in low density lipid rafts (fractions 9-14) when compared to wildtype cells (blue line).

### EhROM1 localizes to lipid rafts

Since membrane composition may influence the activity of rhomboid proteases [34,35], we tested if EhROM1, itself, was compartmentalized in raft-like domains. Western blot analysis of gradient-purified rafts revealed that EhROM1 was, in fact, also enriched in lipid rafts (Fig. 3A). Therefore, like other intramembrane proteases [34,35], EhROM1 may rely on a sterol-rich environment of rafts for its function.

**Fig. 3.**
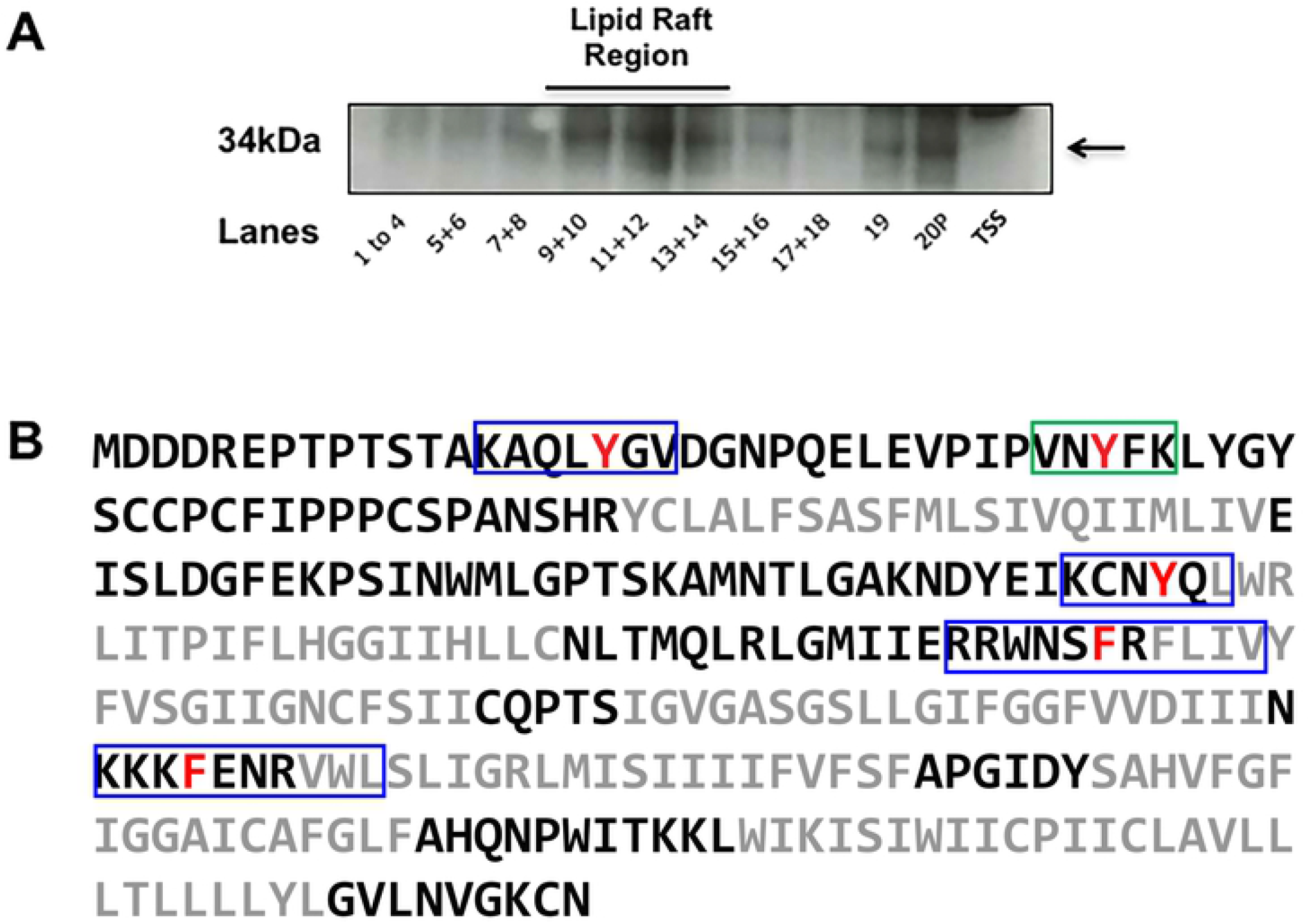
EhROM1 is localized to lipid rafts and possesses putative cholesterol-binding motifs. A. Triton-insoluble membranes were isolated from wildtype cells and resolved by sucrose gradient density centrifugation. Nineteen fractions and the non-buoyant pellet (20P) were collected and subjected to western blot analyses using antibodies specific for EhROM1. TSS represents the triton soluble supernatant. Fractions 1 through 4, and pairs of fractions thereafter, were combined prior to analysis. EhROM1 is enriched in fractions 9 through 14, which have been shown previously to contain lipid rafts [18]. EhROM1 could not be detected in the TSS. B. The amino acid sequence of EhROM1 in which putative CRAC (green box) and CARC sequences (blue boxes) are indicated. The central conserved tyrosine or phenylalanine of each of the sequences is highlighted in red. EhROM1 has 7 putative membrane domains, shown in gray font. These transmembrane regions are predicted by Uniprot (Uniprot KB-C4LVG2 (C4LVG2_ENTH1) [55]

The enrichment of EhROM1 in lipid rafts raised the question of whether the protease can bind directly to cholesterol. Two consensus amino acid sequences for cholesterol-binding have been described in multiple cholesterol-binding proteins [40–43], including the human rhomboid protease, RHBDL4 [35]. The first consensus sequence is the “cholesterol recognition amino acid consensus” (CRAC) sequence, which consists of a leucine or valine residue, followed by 1 to 5 other amino acids, a central tyrosine, 1-5 other amino acids, and finally an arginine or lysine residue (L/V-X_(1-5)_-Y-X_(1-5)_-R/K). The second consensus sequence is a “mirror image” of the first consensus sequence and thus, is referred to as CARC (R/K-X_(1-5)_-Y/F-X_(1-5)_-L/V) [40]. There is a central conserved tyrosine in CRAC sequences and a central conserved tyrosine or phenylalanine in CARC sequences. In either case, these residues seems to be essential for cholesterol-binding [42,43]. Analysis of the EhROM1 amino acid sequence revealed one putative CRAC sequence and four putative CARC sequences (Fig. 3B). Three of the four CARC sequences partically overlap the predicted transmembrane regions of the protein. Consequently, EhROM1 may have the capacity to bind cholesterol directly.

### Cell surface levels of the Gal/GalNAc subunits are higher in T-EhROM1-s cells

We have previously shown that enrichment of Gal/GalNAc lectin subunits in rafts correlates with increased galactose-sensitive adhesion [19]. It would follow that T-EhROM1-s cells exhibit increased adhesion to host cells. Contrary to this prediction, it was reported that inhibition of expression of EhROM1 correlates with defects related to reduced adhesion to host cells [32,33]. The purification of rafts by detergent extraction and sucrose gradient centrifugation is carried out with whole cells. Consequently, the protocol does not distinguish between the adhesins in cell surface lipid rafts versus those in intracellular lipid rafts. Despite raft-enrichment, one reason that T-EhROM1-s cells may have impaired adhesion is because less Gal/GalNAc lectin is in cell surface rafts and more Gal/GalNac lectin is trapped in intracellular rafts. Therefore, we quantified the surface levels of Hgl, Lgl and Igl in T-EhROM1-s cells using biotinylation as previously described [19]. Whole cells were exposed to sulfo-NHS-SS-biotin to label surface proteins. Cell lysates were subjected to avidin-agarose affinity chromatography. Equivalent fractions of the starting cell lysate and affinity purified protein were analyzed for the lectin subunits by western blotting. Surprisingly, surface biotinylation and affinity chromatography revealed that there were significantly higher levels of all three subunits on the surface of mutant cells compared to control cells (Fig. 4). Using immunofluorescence microscopy with α-Hgl antibody, Baxt *et al*. [32] concluded that there was no increase in the surface level of Hgl when EhROM1 is silenced. The incongruence of our data with that of Baxt *et al*. [32] is likely a reflection of the different techniques used to quantify cell-surface proteins. We also observed an increase in the level of *intracellular* lectin subunits. These would be found in the flow-through fractions after biotinylation (Fig. 4). These increases were statistically significant for Igl (\**P*<0.05), but not for Hgl and Lgl. These data are consistent with that of Baxt *et al*., [32], who reported an increase, albeit not statistically significant, in the level of Hgl as detected by the ELISA method. Together, with the increases seen for the cell surface-localized subunits in this study, these observations suggest that there may be more Hgl, Lgl, and Igl, overall in the mutant cell line.

**Fig. 4.**
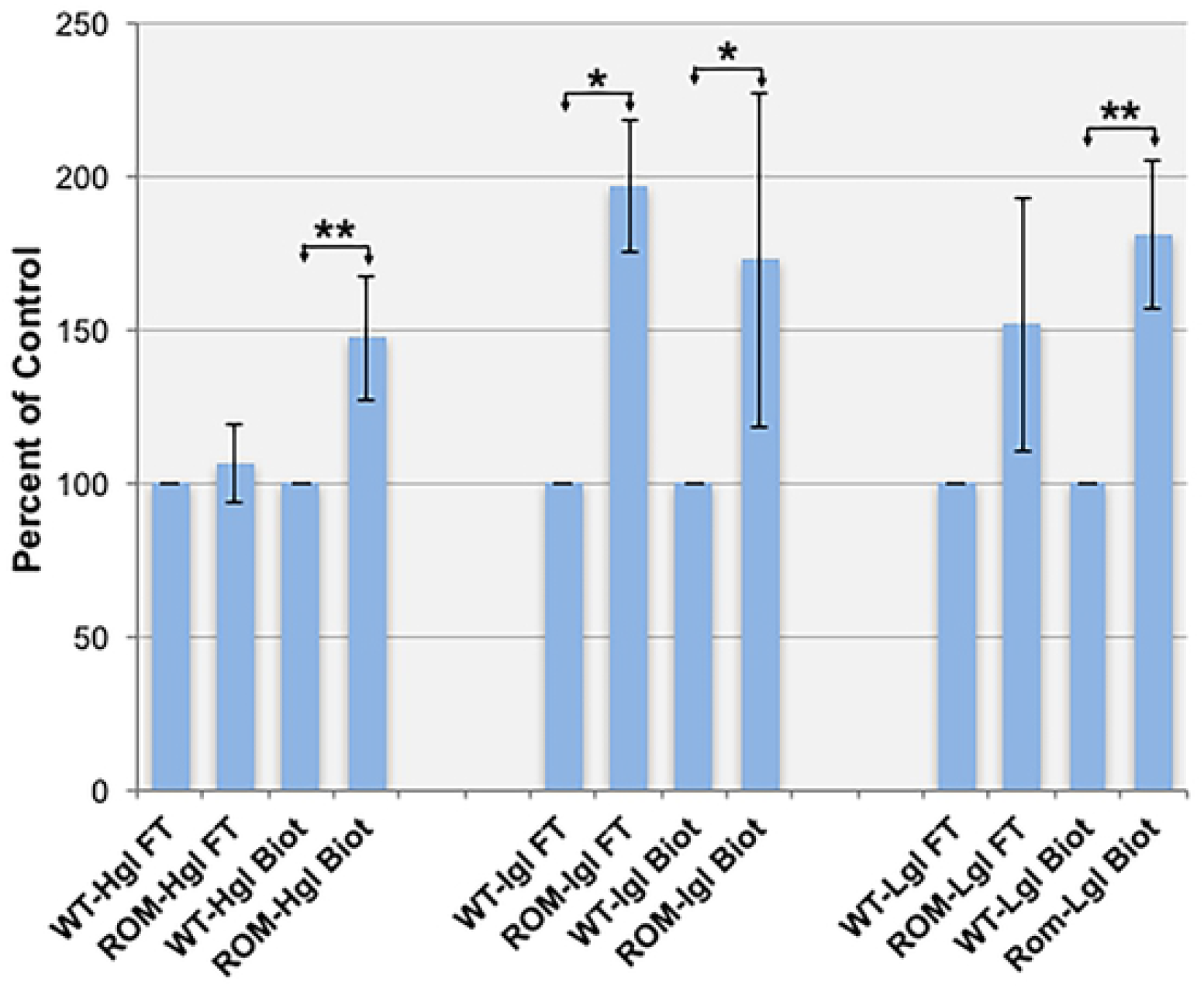
The Gal/GalNAc lectin subunits are enriched on the surface of T-EhROM1-s cells. Wildtype (WT) and T-EhROM1-s (ROM) cells were surface biotinylated. Labeled surface proteins were captured by avidin affinity chromatography. Both purified biotinylated surface proteins (Biot) and the flow through fraction (FT) were analyzed by SDS-PAGE and western blotting with antibodies specific for the heavy subunit (Hgl), the intermediate subunit (Igl) or the light subunit (Lgl). The data represent mean values of densitometric scans (n ≥ 3), reported as a percentage of total Hgl, Igl, or Lgl in WT cells, which was arbitrarily set to 100%. There is a statistically significant increase of all three subunits on the surface of the amoebae (\**P*<0.05; \*\**P*<0.01). There are increases in the intracellular levels of Igl, Lgl, and Hgl as shown by analysis of the flow-through (FT) fractions. Only the increase in the intracellular level of Igl is statistically significant (\**P*<0.05).

### Hgl and Igl exhibit a higher molecular weight in T-EhROM1-s cells

It was intriguing that the T-EhROM1-s cell line exhibited enrichment of the Hgl and Lgl in rafts, and higher levels of the Gal/GalNAc lectin subunits at the surface, but was impaired in many adhesion-related functions [32,33]. Since Hgl is a putative substrate for EhROM1 [31], perhaps proper cleavage of Hgl, by EhROM1 is required for suitable function of the Gal/GalNAc lectin. To test this, we isolated lipid rafts and examined the size of the lectin subunits in both mutant and control cells using SDS-PAGE and western blotting in a MOPS buffer system, which provides superior separation of high molecular weight species. In the mutant cell line, Hgl was larger than its counterpart in control cells (Fig. 5). This was true for this subunit in all membrane domains (i.e., rafts, actin-rich membrane, and detergent-sensitive membrane) (Fig. 5). However, we were surprised to see that the size of Igl was also increased in all membrane domains in the mutant cell line (Fig. 5). These data support the conclusion that Hgl is a substrate of EhROM1 and suggest that Igl is also a substrate of this protease. There is evidence that rhomboid proteases can act on hydrophilic sequences outside of transmembrane domains [44–46]. Thus, it is conceivable that GPI-linked Igl could be a substrate of EhROM1 even though it is predicted to reside completely outside of the membrane.

**Fig. 5.**
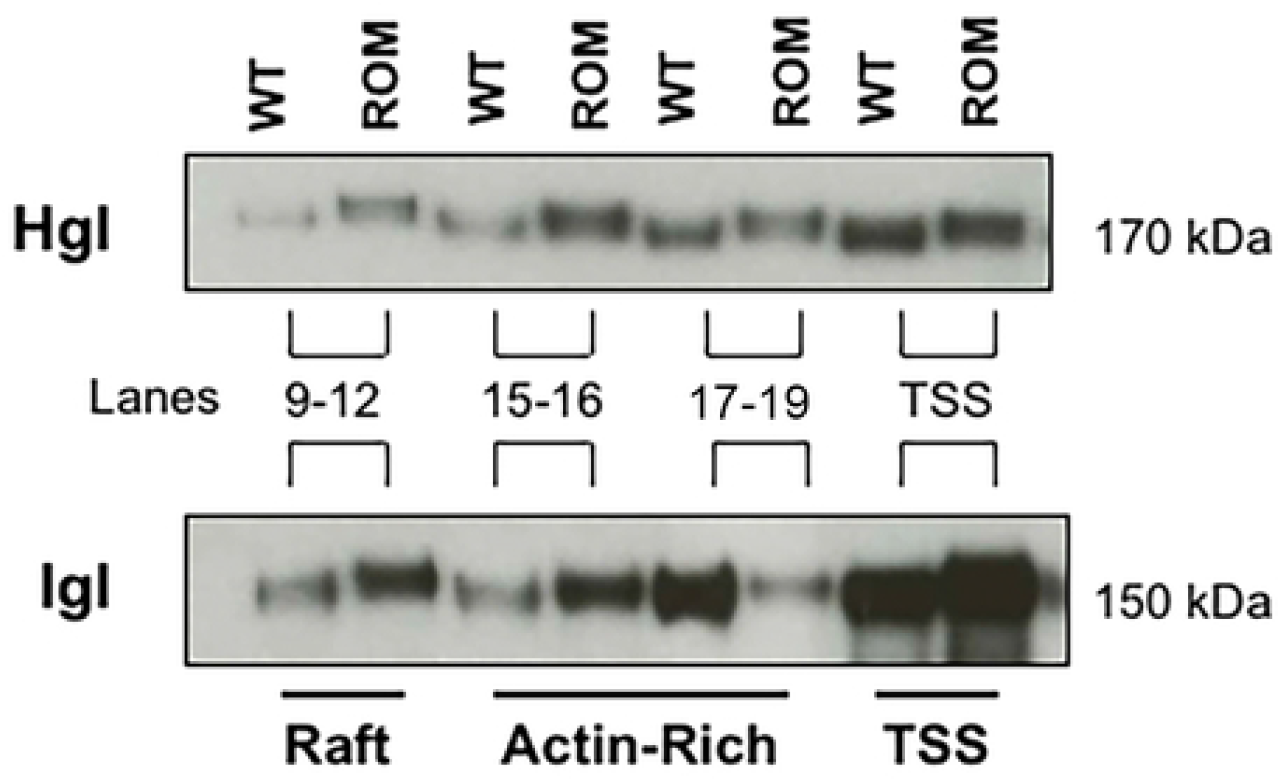
Hgl and Igl exhibit a higher molecular weight in T-EhROM1-s cells. Triton-insoluble membranes were isolated from wildtype (WT) or T-EhROM1-s (ROM) cells and resolved by sucrose gradient density centrifugation. Nineteen fractions were collected. Fractions 9 through 12 (lipid rafts), 15 and 16 (actin-rich membrane), or 17 through 19 (actin-rich membrane) were combined and subjected to SDS-PAGE in a MOPS buffer system, which superior in its ability to resolve high molecular weight proteins. Following SDS-PAGE, western blot analysis was carried out using antibodies specific for the heavy subunit (Hgl), or the intermediate subunit (Igl). The triton-soluble supernatant (TSS) was also analyzed in the same manner. Hgl and Igl are bigger in all membrane types in the mutant cell line.

### Assembly of the Gal/GalNAc lectin trimer is not inhibited in T-EhROM1-s cells

The shift in size of Hgl and Igl in the T-EhROM1-s cell line raised the possibility that the subunits are not interacting with each other normally. This, in turn, could explain the adhesion defect in the mutant cell line. To test this, we examined interaction among the subunits using an immunoprecipitation approach. Detergent-sensitive membrane and various fractions of detergent-resistant membrane, resolved by sucrose gradient centrifugation, were mixed with anti-Hgl monoclonal antibody. Following magnetic purification, the immune complexes were resolved by SDS-PAGE and analyzed by western blotting using α-Hgl, α-Lgl, or α-Igl antibodies. In all membrane types in the mutant cells, there was no alteration in the interaction profiles of the Gal/GalNAc lectin subunits (Fig. 6). Therefore, the increased size of Hgl and Igl in the T-EhROM1-s cell line did not seem to hinder the assembly of trimers, and thus, cannot explain the adhesion defect. Perhaps improper cleavage of the subunits masks crucial binding sites and/or alters the three-dimensional structure of the trimer, in a manner that inhibits its function.

**Fig. 6.**
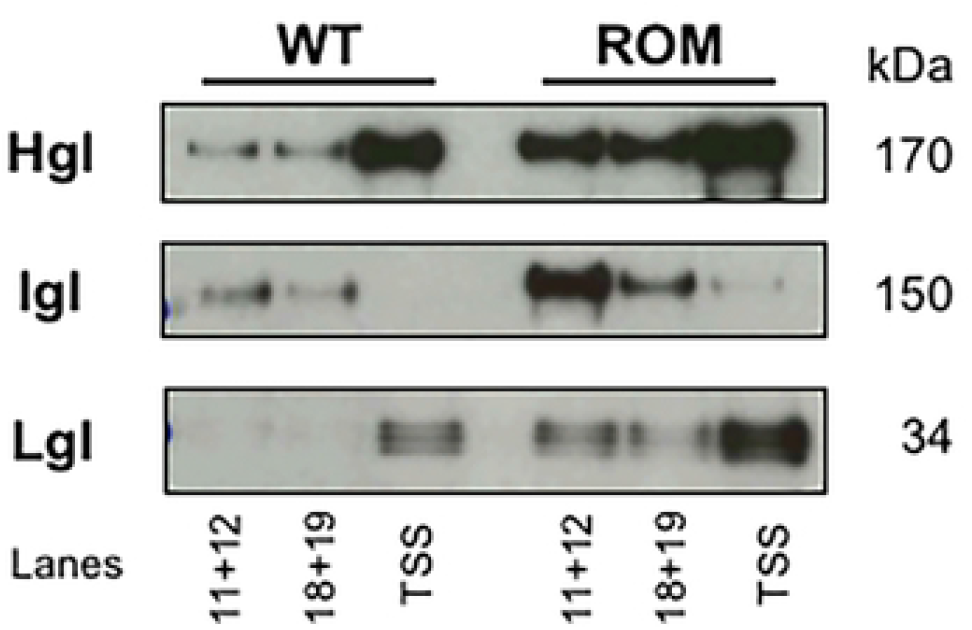
Assembly of the Gal/GalNAc lectin trimer is not inhibited in T-EhROM1-s cells. Triton-insoluble membranes were isolated from wildtype (WT) or T-EhROM1-s (ROM) cells and resolved by sucrose gradient density centrifugation. Nineteen fractions were collected. Fractions 11 and 12 (lipid rafts), or 18 and 19 (actin-rich membrane), were combined and subjected to immunoprecipitation using a monoclonal α-Hgl antibody. The triton-soluble supernatant (TSS) was also analyzed in the same manner. Precipitated proteins were resolved by SDS-PAGE and assessed by western blotting using antibodies specific for the heavy subunit (Hgl) (polyclonal), the intermediate subunit (Igl) or the light subunit (Lgl). Pull down of Hgl reveals co-association with Lgl in all membrane types. In T-EhROM1-s cells, more Lgl associates with Hgl in the lipid rafts (11 +12) because both subunits are enriched in rafts in that cell line. Little Igl co-associates with Hgl in the TSS, because the majority of Igl is constitutively in lipid rafts.

## Discussion

The major finding of this study is that rhomboid protease activity may regulate the subunit size and the sub-membrane location of the Gal/GalNAc lectin in *E. histolytica*. Specifically, we demonstrated that EhROM1, itself, was enriched in rafts. Furthermore, reducing rhomboid protease activity, either pharmacologically or genetically, correlated with an enrichment of Hgl and Lgl in rafts. In the mutant cell line with reduced EhROM1 expression, there was also a significant augmentation of the level of all three Gal/GalNAc subunits on the cell surface and an increase in the molecular weight of two of the subunits, Hgl and Lgl.

In previous studies, enrichment of Hgl and Lgl in lipid rafts was correlated with an increase in galactose-sensitive adhesion to host components [19]. However, in the current study, we observed enrichment of Hgl and Lgl in lipid rafts in a cell line known to display defects related to adhesion [32,33]. Thus, for the first time, we show that enrichment of the Hgl-Lgl dimer in lipid rafts is not sufficient to enhance the adhesive capacity of the parasite. One possible explanation for the accumulation of Hgl and Lgl in rafts in the mutant cell line is that cleavage of the Gal/GalNAc lectin subunits by EhROM1 is necessary for recycling or exit of these subunits out of rafts. This would be consistent with previous reports that demonstrate that rhomboids play a role in dismantling adhesion complexes in a variety of parasites (reviewed in [20]) including *Plasmodium* [27,28,47,48], *Toxoplasma* [22], and *Trichomonas* [30]. Here, we also see, for the first time, an enrichment of the Hgl-Lgl dimer in lipid rafts in the absence of extracellular ligand binding. This insinuates a model in which the dimer is constantly entering and exiting rafts, with the majority of Hgl-Lgl outside of rafts at equilibrium. If egress from rafts relies on rhomboid protease activity, then a lack of rhomboid expression would naturally alter this balance resulting in an accumulation of the dimer in these microdomains.

That the lectin subunits require EhROM1 activity to exit rafts would also be consistent with a hypothesis put forth by Baxt *et al*. [31]. Specifically, these authors demonstrated that EhROM1 was localized to caps, also known as uroids, which are structures that are also rich in Gal/GalNAc lectin and found at the trailing edge of motile cells. Uroids form during motility for the purpose of concentrating and shedding bound host proteins, such as antibodies. Based on these findings, Baxt *et al*. [31] hypothesized that EhROM1 regulates the release of individual lectin subunits from uroids. This proposition is supported by the current study, which shows that EhROM1 is enriched in rafts and that the Hgl-Lgl dimer accumulates in rafts in cells with reduced EhROM1 expression. This is further supported by our previous study, which shows that uroids are, themselves, rich in rafts [49].

Riestra *et al*. [30] mutated the conserved rhomboid binding site of a substrate of *T. vaginalis* ROM (TvROM), which rendered that substrate unavailable for cleavage. Expression of this mutated substrate resulted in the accumulation of that substrate on the cell surface and increased *T. vaginalis*-host adhesion. We also observed a significant increase in the level of the Gal/GalNAc lectin subunits on the surface of the cell line with reduced EhROM1 expression. Remarkably though, this increase was *not* correlated with enhanced adhesion. In fact, lower EhROM1 expression is correlated with reduced adhesion [32,33]. Perhaps the adhesion-related defects in the mutant cell line are a result of the aberrant size of Hgl and Igl. Immunoprecipitation assays demonstrated that the larger Gal/GalNAc subunits were able to interact with each other in a normal fashion; but, complexes possessing these larger subunits may no longer be able to interact with host ligands and/or other *E. histolytica* proteins necessary for adhesion. For instance, crucial binding sites may be hidden and/or the three-dimensional structure may be aberrant in un-cleaved lectin subunits.

The increased molecular weight of the intermediate subunit in the mutant cell line is particularly intriguing. First, it suggests that Igl is a substrate of EhROM1, even though this subunit is surmised to be GPI-anchored and localized completely *outside* the membrane. More recently it has been shown that some rhomboid proteases can cleave sequences within hydrophilic sequences outside of transmembrane domains [44–46]. In these cases, the hydrophilic peptide substrates bend into the protease active site from above the membrane plane (reviewed in [50]). Our observations suggest the EhROM1 is also capable of cleaving sequences outside of the membrane environment.

The increased size of Igl in the mutant cell line is also interesting because unlike Hgl, Igl does not possess an obvious conserved rhomboid protease cleavage site. Indeed, it has already been established that EhROM1 displays atypical substrate specificity. Along with *Trichomonas vaginalis* ROM1 (TvROM1) [30] and *Plasmodium falciparum* Rhomboid 4 (PfROM4) [27], EhROM1 is one of three parasite rhomboid proteases that cannot process Spitz, a substrate from *Drosophila* that is cleaved by most known rhomboid proteases [31]. Although having a conserved rhomboid substrate amino acid motif would strengthen the idea that Igl is an EhROM1 substrate, there are examples of other authentic rhomboid substrates that lack this conserved sequence. Accordingly, other mechanisms for rhomboid-substrate recognition must exist [50]. For instance, cleavage of thrombomodulin by vertebrate rhomboid-2, RHBDL2, is directed by its cytoplasmic tail rather than by sequences in its transmembrane region [51]. Furthermore, a recent study of the activity of ten diverse rhomboid proteases (1 prokaryotic and 9 eukaryotic), in an *in vitro* reconstituted membrane system, demonstrates that the requirement for specific sequences in substrates is much less stringent than previously thought and instead hydrolysis is likely driven, in part, by substrate concentration (rate-driven) [52]. Thus, co-localization of Igl and EhROM1 in rafts may be sufficient for Igl to be hydrolyzed by EhROM1. In fact, the environment of the lipid raft, itself, may contribute to the ability of EhROM1 to target Igl or other substrates that lack the canonical cleavage signal. Two previous studies [53,54] demonstrated that it was possible to transmute non-substrates into substrates simply by changing membrane composition with membrane-disrupting agents, including the cholesterol-binding agent, methyl-β-cyclodextrin.

Finally, we cannot rule out the possibility that the observed changes in the mutant cell line are not directly related to the loss of EhROM1 activity. Perhaps, reduced cleavage of another unknown substrate indirectly leads to the accumulation of the Hgl-Lgl dimer in rafts and to the change in size of Hgl and Igl. Nevertheless, the study provides significant insight into parasite-host adhesion for *E. histolytica*. Given the importance of adhesion for virulence of *E. histolytica*, new information on the molecular mechanisms governing host colonization may contribute to the development of new drugs for this devastating global disease.

## Acknowledgements

The authors thank Dr. Upinder Singh (Division of Infectious Diseases, Dept. of Internal Medicine, Dept. of Microbiology and Immunology, Stanford University School of Medicine, Stanford, CA, USA) for the T-EhROM1-s cell line and for α-EhROM1 antibodies. The authors thank Dr. William A. Petri, Jr. (Division of Infectious Diseases and International Health, University of Virginia, Charlottesville, VA, USA) for monoclonal α-Hgl, α-Igl, and α-Lgl antibodies.

This work was supported by a National Institute of General Medical Sciences Grant: GM109094 (LAT) and a National Institute of Allergy and Infectious Diseases Grant: AI108287 (LAT). The funding agency had no role in study design, data collection and analysis, decision to publish, or preparation of the manuscript. This content is solely the responsibility of the authors and does not necessarily represent the views of the National Institute of General Medical Sciences, National Institute of Allergy and Infectious Diseases, or the National Institutes of Health.

## Author Contributions

**Conceptualization:** Brenda H. Welter, Lesly A. Temesvari

**Data curation:** Brenda H. Welter

**Formal analysis:** Brenda H. Welter

**Funding acquisition:** Lesly A. Temesvari

**Investigation:** Brenda H. Welter

**Methodology:** Brenda H. Welter, Lesly A. Temesvari

**Project administration:** Lesly A. Temesvari

**Resources:** Lesly A. Temesvari

**Software:** N/A

**Supervision:** Lesly A. Temesvari

**Validation:** Brenda H. Welter

**Visualization:** Brenda H. Welter, Lesly A. Temesvari

**Writing – original draft:** Lesly A. Temesvari

**Writing – review & editing:** Lesly A. Temesvari, Brenda H. Welter

